# Genetic dominance governs the evolution and spread of mobile genetic elements in bacteria

**DOI:** 10.1101/863472

**Authors:** Jerónimo Rodríguez-Beltrán, Vidar Sørum, Macarena Toll-Riera, Carmen de la Vega, Rafael Peña-Miller, Alvaro San Millán

## Abstract

Mobile genetic elements (MGEs), such as plasmids, promote bacterial evolution through horizontal gene transfer (HGT). However, the rules governing the repertoire of traits encoded on MGEs remain unclear. In this study, we uncovered the central role of genetic dominance shaping genetic cargo in MGEs, using antibiotic resistance as a model system. MGEs are typically present in more than one copy per host bacterium and, as a consequence, genetic dominance favors the fixation of dominant mutations over recessive ones. Moreover, genetic dominance also determines the phenotypic effects of horizontally acquired MGE-encoded genes, silencing recessive alleles if the recipient bacterium already carries a wild-type copy of the gene. The combination of these two effects governs the catalogue of genes encoded on MGEs, dictating bacterial evolution through HGT.

## Introduction

HGT between bacteria is largely mediated by specialized MGEs, such as plasmids and bacteriophages, which provide an important source of genetic diversity and play a fundamental role in bacterial ecology and evolution (*1*). The repertoire of accessory genes encoded on MGEs and their ability to be phenotypically expressed in different genetic backgrounds are key aspects of MGE-mediated evolution. There are several factors known to impact horizontal gene transferability in bacteria, such as the level of gene expression, the degree of protein connectivity, or the biochemical properties of proteins (*2–5*), but the specific parameters that shape the repertoire of genes encoded on MGEs remain largely unknown (*6, 7*).

Genetic dominance is the relationship between alleles of the same gene in which one allele (dominant) masks the phenotypic contribution of a second allele (recessive). In diploid or polyploid organisms, dominant alleles stem the establishment of new traits encoded by recessive mutations (an effect known as Haldane’s sieve (*8, 9*)). Most bacteria of human interest carry a single copy of their chromosome. In haploid organisms like these, new alleles are able to produce a phenotype regardless of the degree of genetic dominance of the underlying mutations. Therefore, the role of genetic dominance in bacterial evolution has generally been overlooked. However, the bacterial genome consists of more than the single chromosome; a myriad of mobile genetic elements populate bacterial cells. Many MGEs, including plasmids and filamentous phages, replicate independently of the bacterial chromosome and are generally present at more than one copy per cell, with copy number ranging from a handful to several hundred (*10, 11*). Extra-chromosomal MGEs thus produce an island of local polyploidy in the bacterial genome (*12, 13*). Moreover, HGT in bacteria mostly occurs between close relatives (*14, 15*), and genes encoded on mobile elements can therefore create allelic redundancy with chromosomal genes. In light of these evidences, genetic dominance should strongly affect both the emergence of new mutations in MGE-encoded genes and the phenotypic effects of horizontally transferred alleles.

## Results and Discussion

To test whether genetic dominance determines the emergence of mutations in MGE-encoded genes, we used a two-gene synthetic system that can confer tetracycline resistance through dominant and recessive mutations (Figure 1). This construct consists of a *cI* gene, encoding the bacteriophage λ CI repressor, in control of the expression of a contiguous *tetA* gene, which encodes a tetracycline efflux pump (*16*). This system provides tetracycline resistance when *tetA* transcription is derepressed. Derepression can be achieved through mutations that either inactivate the *cI* gene or disrupt the CI binding site upstream of *tetA* (Figure 1A). The repressor gene provides a large target (714 bp), but we hypothesized that mutations inactivating *cI* would be recessive, since *in-trans* copies of wild type CI would repress *tetA*. In contrast, the CI binding site is a short target (166 bp), but mutations in this region are likely to be dominant because they will lead to non-repressible, constitutive TetA production.

**Figure 1.**
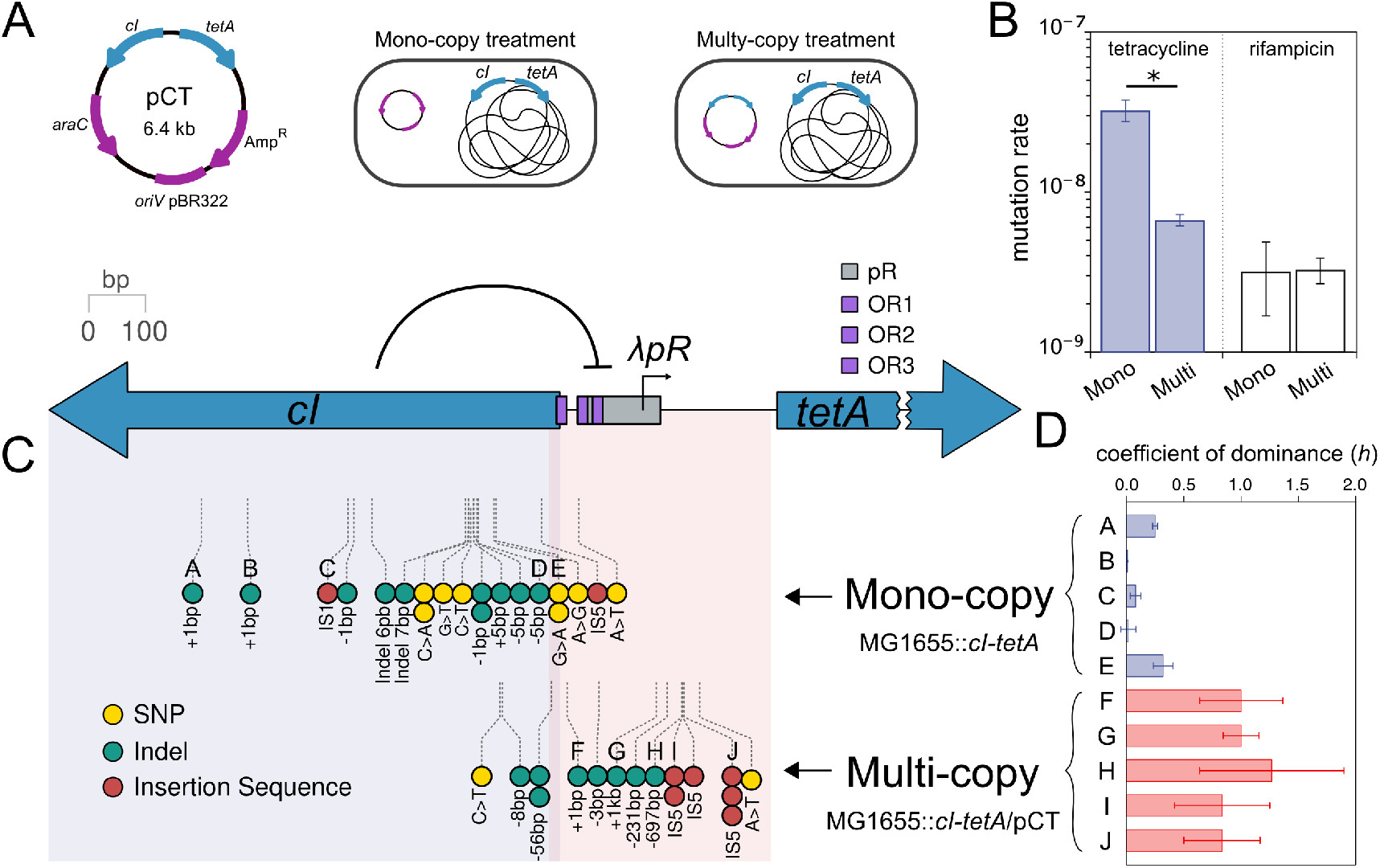
Genetic dominance and gene copy number modulate phenotypic mutation rates. (A) Schematic representation of plasmid pCT, the experimental model and the *cI-tetA* system (note the isogenic nature of the clones apart from the dosage of *cI-tetA*). (B) Phenotypic tetracycline resistance mutation rates in the different clones. Rifampicin mutation rates were determined to test the equal underlying mutation rate of clones. Error bars represent 84% confidence intervals. (C) Location and type of tetracycline resistance mutations in the mono-copy and multi-copy treatments are indicated. In the *cI-tetA* system diagram, blue shading denotes the *cI* coding region, and purple shading denotes the CI binding site plus the *cI-tetA* intergenic region. (D) Coefficient of dominance (*h*) of 10 mutations described in panel C. Bars represent the median of 8 biological replicates, error bars represent the interquartile range.

We produced two otherwise isogenic *Escherichia coli* MG1655 clones carrying the *cI-tetA* system either as a single chromosomal copy (mono-copy treatment) or also present on a pBAD plasmid with approximately 20 copies per cell (pCT, multi-copy treatment) (Figure 1A). Fluctuation assays with both clones revealed a 4.84-fold lower phenotypic tetracycline mutation rate in the multi-copy treatment clone, despite the higher *cI-tetA* copy number (Likelihood ratio test statistic 55.0, *P*< 10^−12^, Figure 1B). To determine if this effect was due to differential access to dominant or recessive mutations, we first analyzed the mutations in the *cI-tetA* system and confirmed that they were located in different regions in each treatment (Wilcoxon signed-rank test, *W*= 313, *P*≈ 10^−06^, Figure 1C, Table S1). Next, we generated homozygous and heterozygous mutant clones in order to measure the coefficient of dominance (h) of a subset of mutations [h ranges from 0 (completely recessive) to 1 (completely dominant), Figure S1–S2, Table S2]. As predicted, *h* was low or intermediate in tetracycline resistance mutations recovered from the mono-copy treatment and high in mutations from the multi-copy treatment (ANOVA effect of treatment; F=70.02, df=1, P≈3×10^−5^; Figure 1D).

Our results indicate that interplay between genetic dominance and gene copy number determines the rate at which phenotypic mutants emerge in bacteria. The increased gene dosage provided by MGEs improves the chances of a beneficial mutation being acquired but simultaneously masks the phenotypic contribution of the newly acquired allele if it is recessive. To study the general effect of this interplay on the evolution of MGE-encoded genes, we developed a computational model based on the classic fluctuation assay ((*17*), Figure 2A and S3–S4). This model allows us to simulate the acquisition and segregation of mutations located in an extra-chromosomal MGE, in this case a plasmid, with a given copy number. With this information, we can explore the frequency of phenotypic mutants in the bacterial population at any time point; this frequency will depend on the distribution of mutated and wild-type alleles in each individual cell and on the coefficient of dominance of those mutations (Figure S4). The simulations showed that the frequency of phenotypic mutants increases with plasmid copy number for mutations of high dominance but decreases for mutations of low dominance (Figure 2A).

**Figure 2.**
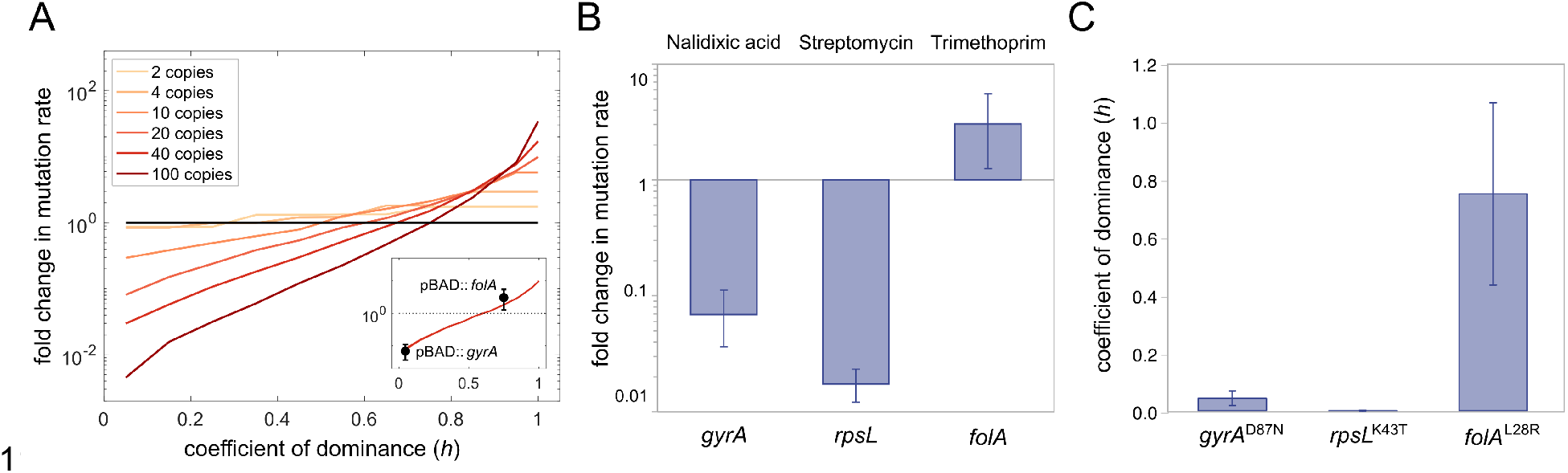
Interplay between genetic dominance and gene copy number. (A) Results of simulations analyzing the effect of plasmid copy number and genetic dominance of a mutation on the emergence of phenotypic mutants. The chart shows fold-changes in phenotypic mutation rate for a plasmid-carried gene at different copy numbers compared with a chromosomal copy of the same gene (black line). The insert chart shows the comparison of experimental results obtained for *gyrA* and *folA* (presented in panels B and C; bars, 84% confidence intervals) with the prediction for a plasmid of 20 copies. (B) Fold change of antibiotic resistance phenotypic mutation rates in *E. coli*, comparing multi-copy and mono-copy treatments for *gyrA, rpsL* and *folA*. Note that result for *rpsL* is an upper bound due to the absence of phenotypic mutants in the multi-copy treatment (see methods). (C) Coefficient of dominance of *gyrA*^D87N^, *rpsL*^K43T^ and *folA*^L28R^. Bars represent the median of 8 biological replicates, error bars represent the interquartile range.

The results of the simulation prompted us to test our hypothesis in a more realistic and meaningful experimental system. To study the effect of genetic dominance and gene copy number on the emergence of phenotypic mutants, we used bacterial housekeeping essential genes known to confer antibiotic resistance through mutations. The genes studied were *gyrA* (DNA gyrase subunit A), *rpsL* (30S ribosomal protein S12), and *folA* (dihydrofolate reductase). Mutations in these genes, which are present as single copies in the chromosome, confer resistance to quinolone, aminoglycoside and trimethoprim antibiotics, respectively (*18, 19*). For each gene, we produced two otherwise isogenic *E. coli* MG1655 clones with either one copy of the gene (chromosomal) or multiple copies (chromosomal + plasmid, Figure S5). We then calculated phenotypic mutation rates for each clone using fluctuation assays with the appropriate antibiotics and sequenced the target genes in the resistant clones to confirm the presence of mutations (Figure 2B, Table S1). The frequency of *gyrA* and *rpsL* mutants in the multi-copy treatment was lower than in the mono-copy treatment (Likelihood ratio test statistic 49.48, *P*<10^−12^), suggesting that antibiotic resistance mutations in these genes are recessive, in line with previous works in the field (*20, 21*). Conversely, the mutation rate for *folA* in the multi-copy treatment increased 3-fold (Likelihood ratio test statistic 5.38, P= 0.04), suggesting that this gene confers trimethoprim resistance through dominant mutations. To test this possibility, we determined the coefficient of dominance of a common mutation in each gene, confirming that mutations in GyrA^D87N^ (h=0.043) and RpsL^K43T^ (h=0) were recessive, whereas mutation in FolA^L28R^ (h=0.748) was dominant (Figure 2C, S1 and S6, for more details on the mechanistic basis of genetic dominance of resistance mutations see supplementary text).

While these results show that genetic dominance shapes the emergence of mutations in MGE-encoded genes, they leave open the question of how genetic dominance affects the horizontal transferability of genes in bacteria. We hypothesized that phenotypic expression of recessive alleles encoded on MGEs will be masked if the recipient bacterium possesses a chromosomal copy of the dominant allele, effectively hindering transfer of the recessive allele. We tested this hypothesis in an experimental assay of bacterial conjugation, using the low copy-number mobilizable plasmid pSEVA121 (*22*). The plasmid was modified by independent insertion of mutated, resistance-conferring *gyrA*^D87N^ (recessive) and *folA*^L28R^ (dominant) alleles and their wild-type counterparts (Figure S7). We transferred these plasmids from *E. coli* ß3914 to *E. coli* MG1655 (Figure 3A). After conjugation, resistance conferred by folA^L28R^ was readily expressed in the recipient cells, whereas resistance conferred by gyrA^D87N^ was masked by the resident wild-type allele, preventing phenotypic expression of resistance in transconjugants (Figure 3A). A simple model of the general effect of genetic dominance on the transferability of housekeeping genes predicted that the presence of a dominant allele in the recipient cell would reduce phenotypic expression of recessive alleles over a wide range of plasmid copy numbers (Figure 3B). Crucially, this effect will be particularly marked for conjugative plasmids, due to their low-copy number in the host cell (typically ranging from 1 to 5).

**Figure 3.**
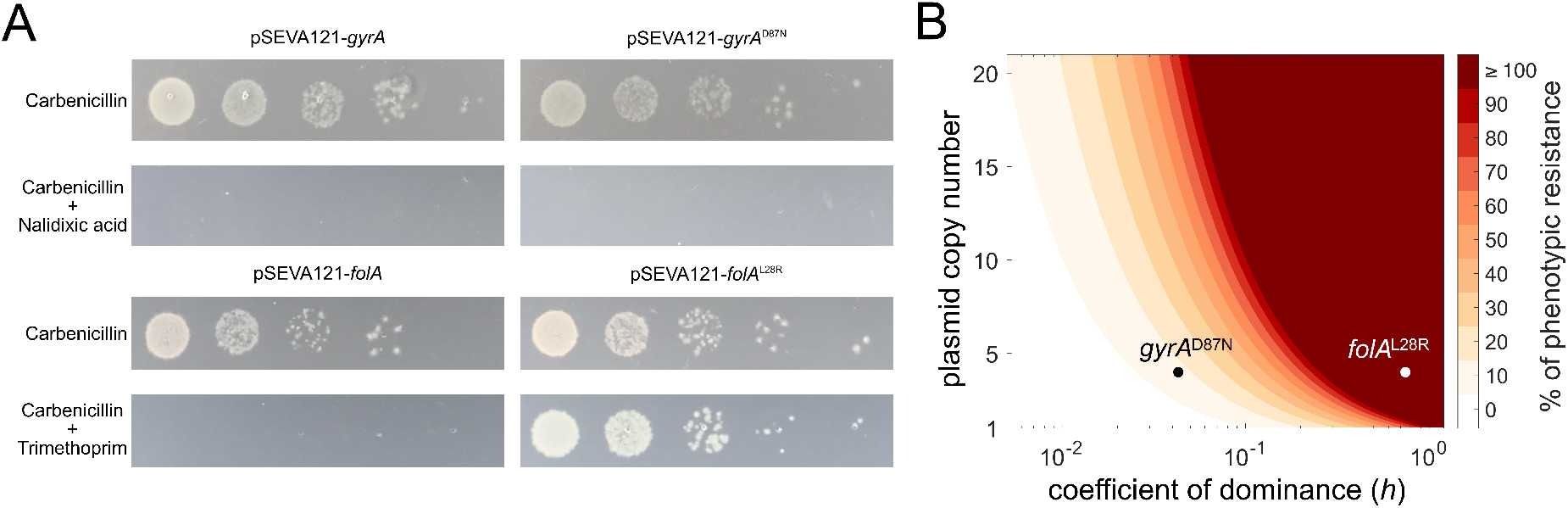
Genetic dominance limits the phenotypic contribution of horizontally transferred recessive alleles. (A) Pictures of a representative replicate of the conjugation assays. Overnight cultures of spots inoculated from 10-fold dilutions of conjugation mixes (10^0^ to 10^−4^, from left to right) on plates selecting for transconjugants. Selection on carbenicillin reveals the actual number of transconjugants; selection on carbenicillin plus nalidixic acid or trimethoprim reveals the number of transconjugants carrying *gyrA* and *folA* alleles and expressing the resistant phenotype. (B) Antibiotic resistance level conferred by a plasmid-encoded resistant allele in the recipient bacterium (when a wild-type copy of the gene is present in the chromosome), assuming phenotypic resistance as the product of plasmid copy number and the coefficient of dominance of the allele. Experimental data are presented for gyrA^D87N^ and *folA*^L28R^ in plasmid pSEVA121.

Our results strongly suggest that genetic dominance shapes the repertoire of genes present in MGEs in bacteria. To investigate this further, we analyzed the Comprehensive Antibiotic Resistance Database (CARD), which includes detailed information about antibiotic resistance genes from thousands of bacterial chromosomes and plasmids (*23*). To extend our analysis to other MGEs, we also examined databases for information on integrative and conjugative elements and prophages. We predicted that housekeeping alleles conferring antibiotic resistance and contained in MGEs would be more frequently dominant than recessive. We investigated the genes in our experimental system plus two additional housekeeping genes that, according to results from previous works, should confer antibiotic resistance through dominant (*folP*, dihydropteroate synthase, sulfonamide resistance) and recessive (*rpoB*, RNA polymerase subunit beta, rifampicin resistance) mutations ((*24–26*), see supplementary text). Results confirmed that mobile resistance-conferring alleles of *folA* and *folP* were ubiquitous in naturally occurring MGEs, whereas resistant alleles of *rpoB, gyrA* and *rpsL* were almost never present in MGEs (Figure 4). Alternative explanations to genetic dominance, such as protein connectivity or the fitness effects produced by these genes in recipient bacteria, could also explain the observed bias in gene distribution. However, analysis of protein connectivity and direct measurements of growth rates did not offer a satisfactory explanation for our results (Figure S8–S9).

**Figure 4.**
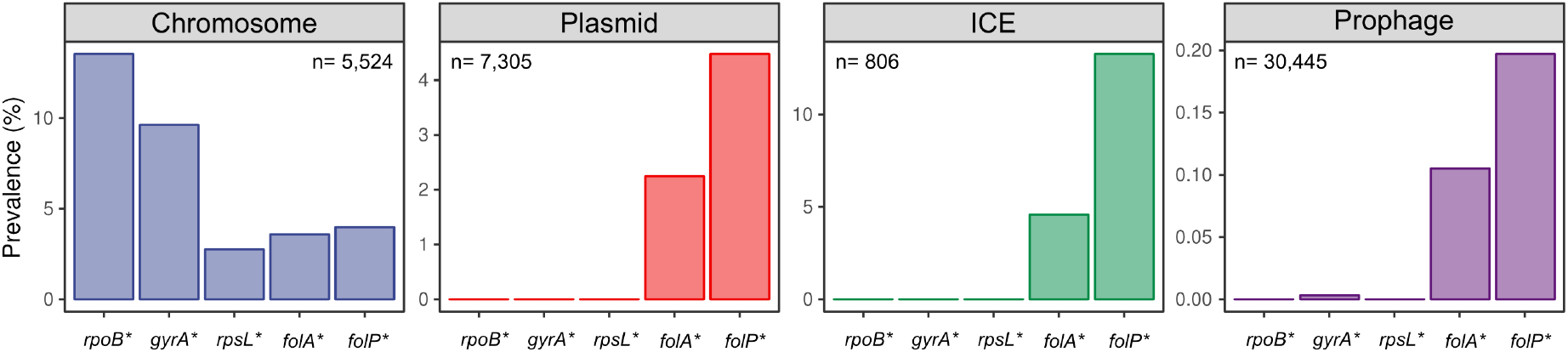
Genetic dominance shapes the bacterial mobilome. Distribution of antibiotic-resistance conferring alleles of *rpoB, rpsL, gyrA, folA* and *folP* (indicated with an asterisk) in chromosomes and MGEs across bacteria. Note that recessive alleles are absent from MGEs. For a more exhaustive analysis see Figure S10 and supplementary text.

In light of these results, we propose that genetic dominance provides a general explanation for the commonly observed divergence between chromosomal-mediated and MGE-mediated antibiotic resistance genes to, for example, fluoroquinolones or polymyxins (*27, 28*). Our results suggest that chromosomal singletons are free to explore their mutational landscape in a host bacterium, but only dominant alleles are able to provide a selectable phenotype, reach fixation and spread successfully on MGEs. MGEs are therefore enriched in the few dominant resistant alleles of chromosomal genes or in xenogeneic genes with no resident counterpart in the host bacterium.

In summary, here we demonstrate that genetic dominance strongly influences evolution through MGEs in bacteria, revealing a new layer of complexity in the forces governing HGT. The impact of genetic dominance on MGEs is twofold, affecting both (i) the emergence of new mutations in MGE-encoded genes and (ii) the phenotypic effects of horizontally transferred genes. The first effect is driven by the polyploid nature of extra-chromosomal MGEs, which induces the filtering of recessive mutations through Haldane’s sieve. However, elevated gene copy number is not an exclusive property of MGEs and can arise physiologically in haploid bacteria in multiple circumstances. For example, gene copy number is increased by gene duplication events and by transient polyploidy during fast bacterial growth (*29, 30*), suggesting a potential broader influence of genetic dominance on bacterial evolution. The second effect is determined by the dominance relationships that emerge when a new allele arrives in a bacterium that already carries a copy of that gene. Given that HGT in bacteria is strongly favored between close relatives (*14, 15*), genetic redundancy of this type must be extremely common, underlining the importance of genetic dominance in determining gene transferability.

## Materials and Methods

### Strains, plasmids and media

All the strains and plasmids used in this study are detailed in Table S3. Experiments were performed using BBL Muller Hinton II agar or cation adjusted broth (Becton Dickinson) unless specified. Antibiotics were supplied by Sigma-Aldrich and were used at the following concentrations: carbenicillin 100 μg/ml, chloramphenicol 30 μg/ml, nalidixic acid 30 μg/ml, streptomycin 100 μg/ml, rifampicin 100 μg/ml, trimethoprim 2 μg/ml and 16 μg/ml, and tetracycline 15 μg/ml. Cultures were routinely grown at 37°C with shaking (225 rpm). Plasmids were extracted using a commercial miniprep kit (Macherey-Nagel) and were transformed into TSS competent cells (*31*).

### Cloning and site directed mutagenesis

The *gyrA, rpsL* and *folA* wild-type alleles were PCR amplified using HS Taq Mix (PCR biosystems) polymerase, using the primers listed in table S4 and MG1655 chromosomal DNA as template. The cI-*tetA* system was amplified similarly, but using IBDS1 as a template (*16*). Purified PCRs were subsequently cloned into the p_BAD_ TOPO TA Expression kit (Thermo-Fisher) following manufacturer instructions. We selected clones in which the insert was cloned in the opposite direction from that of the PBAD promoter to ensure that expression is driven only from the cognate promoter. Correct clones were identified by PCR and Sanger sequenced to give rise to pBAD-gene and pCT plasmids (Table S3).

Site-directed mutagenesis was performed on the pBAD plasmids carrying wild-type alleles to construct the *gyrA*^D87N^, *rpsL*^K43T^ and *folA*^L28R^ mutated alleles by using the Q5 site-directed mutagenesis kit (New England Biolabs) and the primers listed in table S4. Correct cloning was assessed by PCR and Sanger sequencing. Next, we generated pBDSM plasmid variants by Gibson assembling PCR amplified (Phusion Hot Start II DNA Polymerase, Thermo-Fisher; see Table S4 for primers) mutated and wild-type alleles into the pBGC backbone (Figure S5 and Table S3)

Tetracycline resistant mutations in the pCT plasmid isolated in the fluctuation assays were purified by plasmid extraction and re-transformed into MG1655, selecting in carbenicillin and tetracycline plates to isolate pCT mutated plasmids from the wild-type version, that could be coexisting under heteroplasmy. Isolated pCT plasmids and chromosomal mutants carrying *cI-tetA* mutated cassette were PCR amplified (Phusion Hot Start II DNA Polymerase; Thermo-Fisher) and simultaneously Gibson assembled (NEBuilder HiFi DNA Assembly kit; New England Biolabs) into the pBAD and pBGC backbones using the primers listed in Table S4 to give rise to pBAD and pBDSM plasmids carrying mutated alleles of the *cI-tetA* system (Figure S1 and Table S3).

### Fluctuation assays for mutation rate determination

Briefly, independent cultures containing carbenicillin were inoculated with ~10^3^ cells and allowed to grow during 18 hours at 37°C with shaking (225 rpm). The following day, appropriate aliquots were plated in antibiotic-containing agar plates to select for spontaneous mutants and in non-selective plates to determine the number of viable bacteria. After incubation at 37°C for 24 hours (48 hours for rifampicin and streptomycin plates) colonies were enumerated. The genes suspected of carrying mutations were PCR amplified and Sanger sequenced using mutant colonies coming from independent cultures as template. Phenotypic mutation rates, defined as the rate at which mutations contributing to the phenotype emerge, 84% confidence intervals and likelihood ratio tests to asses statistical significance were then calculated using the *newton.LD.plating, confint.LD.plating* and *LRT.LD.plating* functions of the Rsalvador package for R (*32*). We plotted 84% confidence intervals because it has been demonstrated that they better convey statistical significance than 95% confidence intervals when dealing with mutation rate data (*33*).

Mono-copy and multi-copy strains of *gyrA* and *rpsL* presented the same antibiotic resistance levels, indicating that target overexpression did not increase the level of resistance for these genes (Table S2). However, and as previously reported (*34*), *folA* overexpression led to a 8-fold increase in the trimethoprim resistance level in the multi-copy treatment. To perform the mutation rate assay, we used 4 times the IC_90_ of trimethoprim for each clone (2 and 16 μg/ml for the mono-copy and multi-copy treatment, respectively). Sequencing of *folA* PCR products revealed mutations in all the colonies tested from the mono-copy treatment, but only for approximately 10% of the ones from the multi-copy treatment. This result could be due to the lack of fixation of *folA* mutations, which can be maintained at a low frequency in heteroplasmy and not be evident in the chromatogram, or due to the access to different resistance mutations in this treatment. To solve this problem and recover only *folA* mutants, we introduced an extra step in the fluctuation assays for the multi-copy treatment, where we restreaked the colonies from 16 to 50 μg/ml trimethoprim plates. This step allowed us to recover colonies carrying *folA* mutations only (avoiding false positives) while at the same time allowing the recovery of colonies carrying *folA* mutations even if the frequency of *folA* mutated alleles in heteroplasmy would have been too low to initially grow on this concentration during a fluctuation assay (avoiding false negatives) (Table S1).

For *rpsL*, we were not able to recover a single resistant colony in the fluctuation assays for the multi-copy treatment (Table S1). Subsequent analyses revealed that the coefficient of dominance of the most common *rpsL* mutation isolated in the chromosome is 0. The complete recesiveness of *rpsL* streptomycin resistance mutations explains the absence of resistant mutants in the multi-copy treatment because, even if the plasmid-mediated copy of *rpsL* mutates and reaches fixation prior to plating, the wild type chromosomal copy of the gene masks the resistance phenotype. Therefore, for phenotypic resistant mutants to appear, both chromosomal and plasmid copies of the gene should mutate (and reach fixation in the cell), which is extremely unlikely. In Figure 2B we present the fold change of streptomycin resistance phenotypic mutation rate using the mutation rate calculated for the monocopy treatment and the limit of detection of the mutation rate for the multi-copy treatment.

Rifampicin mutation rates were performed to confirm that the underlying mutation rate is equal for all clones used in this study (Figure 1 and S11).

### Coefficient of Dominance

The coefficient of dominance (*h*) of mutations ranges from 0, completely recessive, to 1, completely dominant. To calculate *h*, we developed an experimental system based on two compatible plasmids of similar copy number (n≈ 15-20) (*35, 36*), where we cloned the genes of interest carrying the wild type or mutant alleles under investigation (Figure S1). We then transformed *E. coli* MG1655 with the single plasmids and the combinations of plasmids to produce the two possible heterozygous clones. To construct the homozygous mutant clones for *rpsL*^K43T^ and *gyrA*^D87N^, we used the MG1655 background carrying the same mutation in the chromosomal copy of the gene under study. We then performed antibiotic susceptibility assays to determine the resistance phenotypes of the different clones (measured as inhibitory concentration 90, IC_90_, Figure S2 and S6). Using the IC_90_ values we calculated *h* with the following formula:

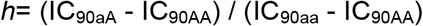

where IC_90AA_, IC_90aA_ and IC_90aa_ are the IC_90_ of the homozygous wild-type, heterozygous and homozygous mutant clones, respectively. Since there are two different heterozygous clones (Figure S1), we calculated *h* as the median value of four independent replicates of each heterozygous clone (n= 8, see Table S2).

IC_90_ values were obtained following (*37*), with some alterations. In short, strains were streaked from freeze stock onto Mueller Hinton II agar plates and incubated over night at 37°C. Single colonies were picked and suspended in liquid Mueller Hinton II broth and incubated at 37°C and 225 rpm. Overnight cultures were diluted 1:10,000 into Mueller Hinton II broth to a final volume of 200 μl per well in microtiter plates. The wells contained increasing concentrations of the appropriate antibiotic in 1.5-fold increments. Plates were incubated at 37°C for 22 hours and 225 rpm. After incubation we ensured homogenous mixture of bacteria by orbital shaking for one minute (548 rpm with 2 mm diameter) before reading of OD_600_ in a Synergy HTX (BioTek) plate reader. IC_90_ is defined as ≥90 % inhibition of growth which was calculated using the formula: 1 – [OD_600_ drug / OD_control_], where the control is the no antibiotic treatment. Wells only containing media were used for background subtraction. Appropriate antibiotic for plasmid maintenance was included in the media throughout the assay.

### Conjugation assays

We developed an experimental model system to determine the frequencies of conjugation of plasmids pSEVA121-gyrA, pSEVA121-gyrA^D87N^, *pSEVA121-folA* and pSEVA121-folA^L28R^ between *E. coli* strains (Figure S7). pSEVA121 is a vector carrying the IncP-type RK2 (also called RP4) replicon, with a *trfA* replication initiator protein gene plus the *oriV* origin of replication, and the *oriT* origin of transfer. pSEVA121 displays therefore low copy-number in the host cell (ca. 4 copies/cell) (*38*), representing a great model system to reproduce the features of a natural conjugative plasmid. To mobilize pSEVA121 we used strain ß3914 as donor, which is auxothropic for diaminopimelic acid and carries RP4 conjugation machinery inserted in the chromosome. As a recipient we used *E. coli* MG1655 (Table S3).

Pre-cultures of donor and recipient strains were incubated overnight in 2 ml of Mueller Hinton broth with the appropriate antibiotics at 37°C and 225 rpm. Next day, 1:100 dilution of the ON cultures in 5 ml of Mueller Hinton broth were incubated until the culture reached mid-exponential phase at 37°C and 225 rpm (2.5 hours approximately). Cultures were then centrifuged for 15 minutes at 1,500 G and the pellets were re-suspended in 200 μl of fresh Mueller Hinton broth. 50 μl of donor and 10 μl of recipient suspensions were mixed and spotted on Mueller Hinton agar plates.

The conjugation mix was incubated for 18 hours at 37°C and then suspended in 2 ml of 0.9% NaCl sterile solution. Dilutions of this suspension were spotted as 5 μl drops on Mueller Hinton agar plates containing antibiotics selecting for donor, recipient and transconjugant cells. The total number of transconjugants was determined by selecting on carbenicillin, while the number of transconjugants expressing the resistance phenotype conferred by *gyrA* and *folA* alleles were selected on carbenicillin plus nalidixic acid or trimethoprim, respectively. We performed 4 independent biological replicates of each conjugation assay, and we show one representative replicate in Figure 3A.

### Growth curves

Growth curves were performed in a Synergy HTX (BioTek) plate reader. Briefly, overnight cultures were diluted 1:1,000 in of Mueller Hinton broth supplemented with carbenicillin, and 200 μl of the dilutions were transferred to a 96 multi-well plate (Thermo Scientific). Plates were incubated at 37°C with strong orbital shaking before reading absorbance at 600 nm every 15 minutes. Six biological replicates were included per strain.

### Bioinformatic analysis

CARD prevalence data containing an analysis of 82 pathogens with more than 89,000 resistomes and more 175,000 antimicrobial resistance allele sequences was obtained from the CARD website (*23*) (downloaded on 18/09/2019).

We downloaded all available bacterial integrative and conjugative elements (ICEs) from ICEberg 2.0 database (*39*), amounting to a total of 806 ICEs from which 662 were conjugative type IV secretion system (T4SS)-type ICEs, 111 were chromosome-borne integrative and mobilizable elements (IMEs), and 33 were *cis*- mobilizable elements (CIMEs). A total of 741 perfect and strict hits from 267 ICEs were identified using CARD RGI (version 4.2.2, parameters used: DNA sequence, perfect and strict hits only, exclude nudge, high quality/coverage, and we used DIAMOND to align the CDS against the CARD database).

We downloaded all complete bacterial genomes from NCBI, a total of 13,169 genomes (Assembly level: complete, downloaded on 15/02/2019). We used the algorithm phiSpy to identify prophages in the bacterial chromosomes and we predicted a total of 30,445 prophages in 7,376 genomes (*40*). To study the presence of genes conferring antibiotic resistance in prophages, we extracted the CDS overlapping with prophages and we predicted their resistome using the command line tool CARD RGI as above.

We then used the antibiotic resistance data from prophages, ICEs, plasmids, and chromosomes to calculate the prevalence of the following Antibiotic Resistance Ontology (ARO) categories: “antibiotic resistant DNA topoisomerase subunit gyrA” (ARO: 3000273), “antibiotic resistant rpsL” (ARO: 3003419), “sulfonamide resistant dihydropteroate synthase” (ARO: 3000558), “antibiotic resistant rpoB” (ARO:3003276), and “antibiotic resistant dihydrofolate reductase” (ARO:3003425).

Connectivity data for all protein families was downloaded from the STRING database (*41*), and the number of high-confidence interactions (>700 STRING confidence) was determined for each protein family.

### Stochastic model

We assume that plasmids replicate randomly throughout the cell cycle until reaching an upper limit determined by the plasmid copy number control mechanism. Therefore, the probability of plasmid replication can be modelled as 1 − *n_i_*(*t*)/*N*, where *n_i_*(*t*) represents the number of copies of a plasmid of type *i* at time *t* and *N* the maximum number of plasmids. We explicitly consider random mutations occurring during replication events, so if *μ* > 0 denotes the probability of a mutation occurring in a plasmid, the per-cell mutation rate is *μ* · *n*. Stochastic simulations of mutation and replication dynamics were performed using a Gillespie algorithm implemented in Matlab with propensities determined from the distribution of plasmid copies of each allele carried by the cell (code can be downloaded from GitHub) (Figure S3).

We also consider that plasmids segregate randomly during cell division. The probability that each plasmid is inherited to one of the daughter cells is a random process that follows a binomial distribution. By implementing an agent-based extension of the replication-mutation model coupled with the segregation dynamics, we simulated the intracellular plasmid dynamics of individual cells in an exponentially growing population (Figure S4). The frequency of phenotypic mutants observed in the population at the end of the numerical experiment was estimated from the fraction of mutated/wild-type alleles in each individual cell and the coefficient of dominance of those mutations (*h*). The results presented in this study were obtained after 10^6^ simulation runs in a range of plasmid copy numbers (Figure 2A). Finally, we used Rsalvador package to estimate from this synthetic data the *-fold* change in phenotypic mutation rate with respect to chromosomally-encoded genes.

### Statistical analysis

All statistical tests were performed using R (v. 3.4.2).

## Acknowledgments

We would like to thank R. Craig MacLean and José R. Penadés for helpful discussion. We are grateful to Ivan Matic, Jesús Blázquez, Jose A Escudero, Didier Mazel and the SEVA collection for generous gifts of strains and plasmids.

## Funding

This work was supported by the European Research Council under the European Union’s Horizon 2020 research and innovation programme (ERC grant agreement no. 757440-PLASREVOLUTION) and by the *Instituto de Salud Carlos III* (grant PI16-00860). ASM is supported by a Miguel Servet Fellowship (MS15-00012). JRB is a recipient of a Juan de la Cierva Fellowship (FJCI-2016-30019). MTR acknowledges support from the Swiss National Science Foundation (Ambizione grant, PZ00P3_161545). RPM was funded by CONACYT Ciencia Básica (grant A1-S-32164). VS is supported by Northern Norway Regional Health Authority (HNF1494-19) and The National Graduate School in Infection Biology and Antimicrobials (NFR grant no. 249062).

## Author contribution

Conceptualization, JRB, ASM. Formal analysis, JRB, VS, MTR, RPM. Funding acquisition, ASM. Investigation, JRB, VS, CdlV. Project administration and supervision, ASM. Software, JRB, MTR, RPM. Writing, JRB, ASM.

## Competing interests

Authors declare no competing interests.

## Data and materials availability

All data is available in the main text or the supplementary materials.

## Supplementary Materials

### Supplementary text

#### Mechanistic basis of genetic dominance of antibiotic resistance mutations analyzed in this work

We experimentally tested the degree of genetic dominance of mutations conferring antibiotic resistance in three housekeeping genes: *gyrA* (GyrA^D87N^), *rpsL* (RpsL^K43T^) and *folA* (FolA^L28R^). Mutations in *gyrA* (DNA gyrase subunit A) and *rpsL* (30S ribosomal protein S12) were recessive, while mutation in *folA* (dihydrofolate reductase) was dominant. The recessive nature of mutations in *gyrA* and *rpsL* can be explained by the toxic effect produced by the combination of antibiotic and wild type alleles. In the case of *gyrA*, when the quinolone antibiotic binds to the gyrase, double-strand DNA breaks occur, leading to cell death (*42*). In the case of streptomycin and *rpsL*, the binding of the antibiotic to RpsL in the ribosome leads to mistranslation events producing a toxic effect in the cell (*43*). Therefore, even if resistance mutations reduce or avoid the binding of the antibiotic to the mutated target, if there are wild type enzymes present in the cell the antibiotic will still produce a toxic effect, explaining the recessive nature of these mutations.

Our experimental results showed that mutations in FolA were strongly dominant. FolA is a key enzyme in folate metabolism. Specifically, it catalyzes the reduction of dihydrofolate to tetrahydrofolate via hydride transfer from NADPH to C6 of the pteridine ring. Trimethoprim binds to FolA and inhibits its activity, blocking folate metabolism and producing a bacteriostatic effect. Mutations in FolA confer trimethoprim resistance through reduction of the binding affinity of FolA for the drug and/or by increasing the activity of this enzyme (both by increasing expression or by specific mutations that increase enzymatic activity (*18, 44*)). However, in this case, the binding between FolA and trimethoprim does not produce a toxic effect *per se*.

Therefore, in the presence of trimethoprim a cell carrying an antibiotic resistant version of the enzyme will be able to grown even if there are wild-type copies of FolA, explaining the genetic dominance of the mutation.

We included two more housekeeping genes known to confer resistance to antibiotics through mutations in our bioinformatic analysis: *rpoB* (RNA polymerase subunit beta, rifampicin resistance) and *folP* (dihydropteroate synthase, sulfonamides resistance, Figure 4). We chose these two genes because the genetic dominance of their resistance mutations could be inferred from previous works. For *rpoB*, the complexes formed between rifampicin and the wild type RNA polymerase block DNA transcription, even if there is a resistant version of RpoB (*24*). Therefore, rifampicin-resistance mutations in *rpoB* are recessive (*24, 25*).

FolP is a dihydropteroate synthase (DPHS) in the folate synthesis pathway and it catalyzes the condensation of para-aminobenzoic acid (PABA) with pteridine diphosphate (PDP) to form dihydropteroate (DHP). Sulfonamides compete with PABA for binding to the enzyme DHPS, resulting in their covalent attachment to PDP in place of PABA (*45*). The resulting product, dihydropterin-sulfonamide, is not toxic to *E. coli* (*46*), and the bacteriostatic activity of these antibiotics comes from the depletion of the essential metabolite PDP. The fact that dihydropterin-sulfonamide is not toxic suggests that *folP* mutations can be dominant (in contrast to *rpsL, gyrA* and *rpoB* mutations). Palmer and Kishony recently described that the fraction of PDP that is converted to DHP (the correct product) or to dihydropterin-sulfonamide (the incorrect one) depends on the ratio of drug to drug-competing substrate (sulfonamide to PABA ratio), weighted by binding affinity of the DHPS enzyme to each substrate (*31*). Resistance mutations in *folP* decrease the binding affinity to sulfonamides drastically (*47, 48*). As a consequence, in this scenario the degree of dominance of mutations is going to depend on the rate of synthesis of DHP by the mutant allele, compared to the rate of synthesis of dihydropterin-sulfonamide by the wild type allele (in other words what fraction of PDP is used in the correct or the incorrect reaction). The rate of synthesis of DHP by the mutant allele depends, in turn, on its binding affinity to PABA. Therefore, the higher the binding affinity of the mutant FolP to PABA the higher coefficient of dominance of the mutation will be. In summary, DHPS resistant alleles will range from mildly dominant (point mutations in *folP* with reduced affinity to PABA (*47*)) to strongly dominant (mobile resistant alleles of DHPS, such as *sul1* and *sul2*, with high binding affinity to PABA (*48*)).

## Supplementary Figures

**Figure S1.**
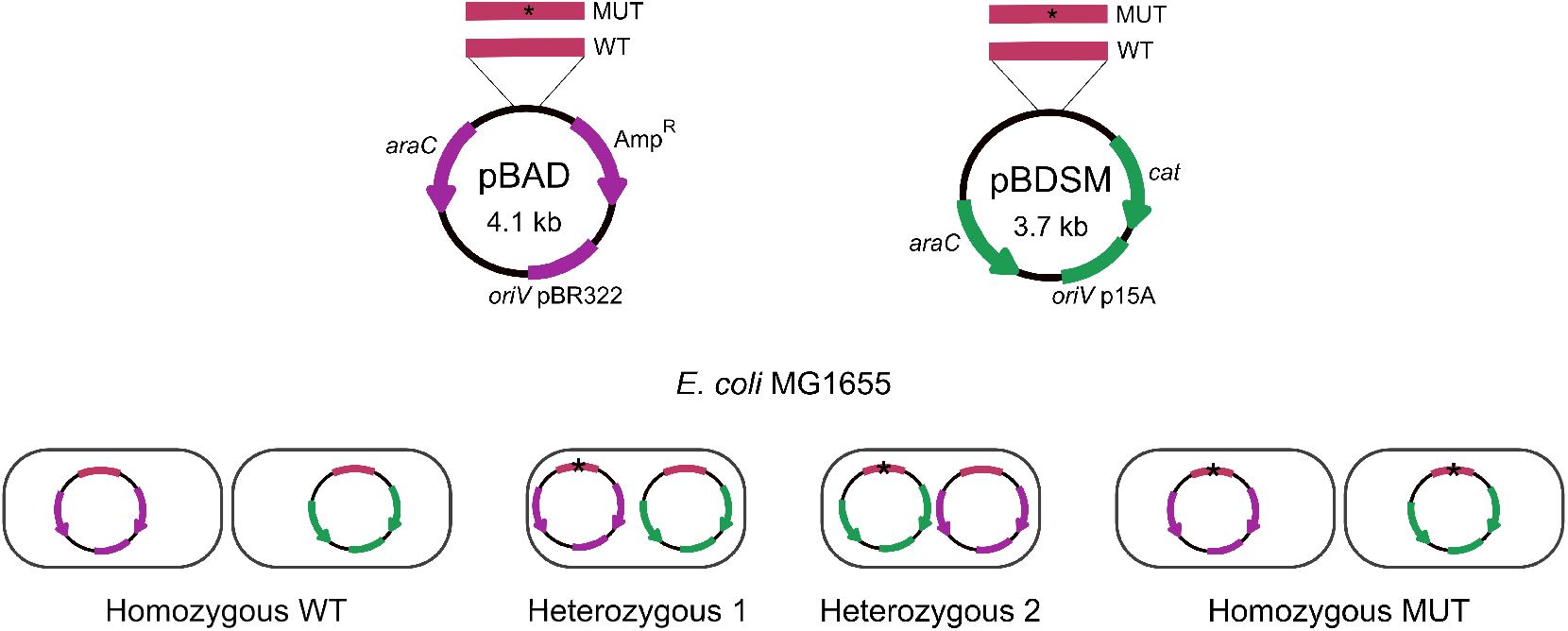
Model system developed to calculate coefficient of dominance of mutations. Schematic representation of plasmids pBAD and pBDSM (Table S1). The gene or genes under study were cloned into these plasmids using Gibson assembly. We cloned both wild type and mutant alleles of the gene or genes under study in both plasmids. We constructed *E. coli* MG1655 strains carrying plasmids creating homozygous wild type and mutant clones plus heterozygous clones (2 different heterozygous combinations per gene or genes under study). To construct the homozygous mutant clones for recessive alleles, we used the MG1655 background carrying the same mutation in the chromosomal copy of the gene under study.

**Figure S2.**
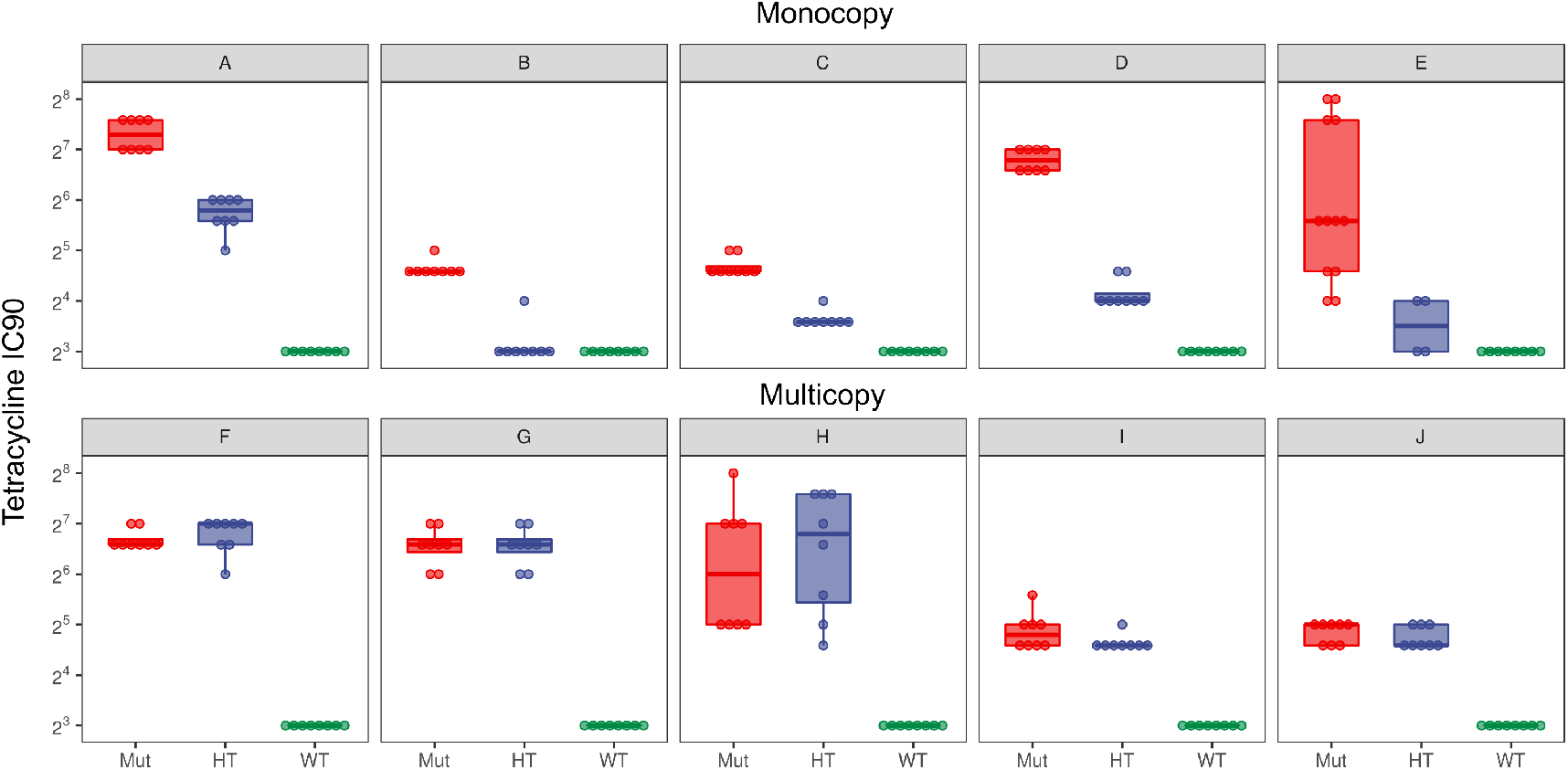
Tetracycline resistance levels of homozygous and heterozygous clones constructed to measure the coefficient of dominance of mutations. Tetracycline resistance phenotypes of the different clones constructed to measure the coefficient of dominance of tetracycline resistance mutations detailed in Figure 1C. The tetracycline inhibitory concentration 90 (IC_90_, in mg/L) of the homozygous mutant clones (Mut), heterozygous mutant clones (HT) and homozygous wild type clones (WT) are represented by boxes. The line inside the box marks the median. The upper and lower hinges correspond to the 25th and 75th percentiles. The letters in the panels correspond to the mutations indicated in Figure 1C. Note that for the mutations isolated in the multicopy treatment the resistance level of HT clones is similar to that of the Mut clones (dominant mutations).

**Figure S3.**
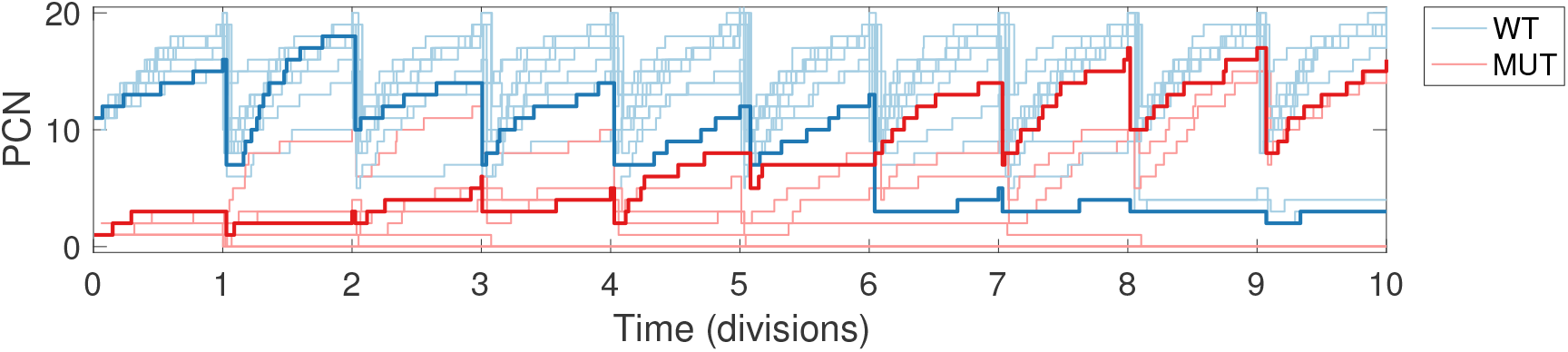
Numerical simulations of the stochastic model of plasmid dynamics in the absence of selection. Replication events increase the number of copies of plasmids carried in a cell until the cell divides and plasmids are segregated randomly between both daughter cells. In this example, we consider that a mutation occurred in a copy of a WT plasmid at t=0 (multiple stochastic realizations of this experiment are shown with light colors): blue lines correspond to the number of copies of the WT plasmid and red lines the number of mutant plasmids as a function of time. Note that in some cases the mutant allele is lost through segregational drift (*49*), while in others (for instance the simulation highlighted with the thick lines) the stochastic nature of the plasmid dynamics produces a gradual increase in mutant frequency.

**Figure S4.**
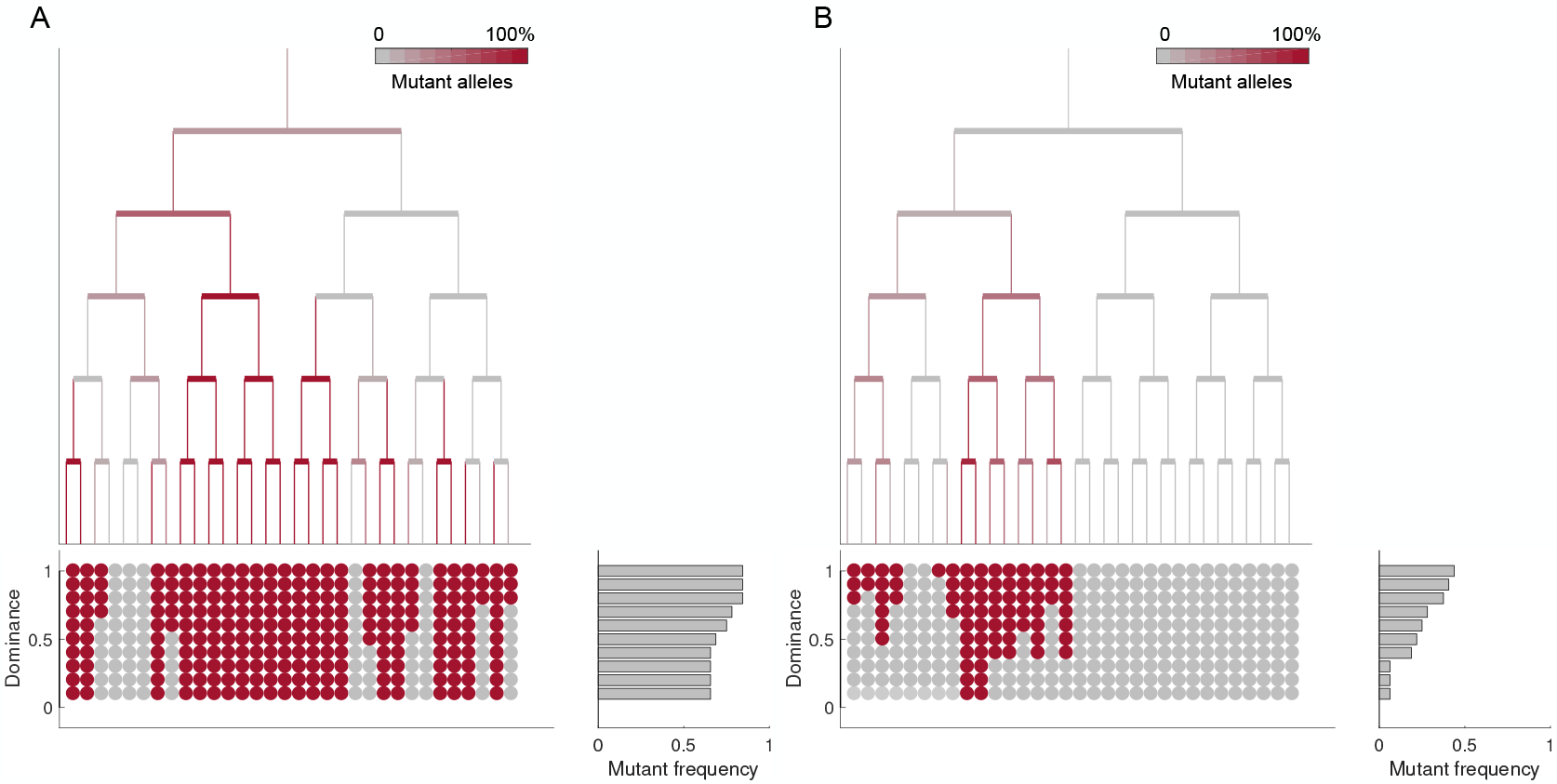
*In silico* fluctuation assays. *In silico* fluctuation assays of populations of cells with a maximum of two plasmids (left) and 15 plasmids per-cell (right). Trees illustrate numerical simulations of the segregation-replication dynamics model in an exponentially growing population of cells in the absence of selection. We consider that mutations can occur randomly during plasmid replication, so we track the fraction of mutant alleles in each cell as a function of time (WT alleles represented in grey, with the fraction of mutant plasmids in each cell is represented with a gradient of red). At the end of the computational experiment, we simulate a fluctuation assay to estimate the phenotypic mutant frequency from the fraction of mutated/wild-type alleles in each individual and the degree of dominance of the mutation. Red circles represent cells that survived the fluctuation assay (bottom). In the case illustrated here, simulations are performed for five generations, but in the synthetic data presented in the manuscript, the experiment was run for 25 generations. Note how, in both cases, the frequency of phenotypic mutants (survivors of the fluctuation assay) decreases as the degree of dominance of the mutation decreases (grey bars).

**Figure S5.**
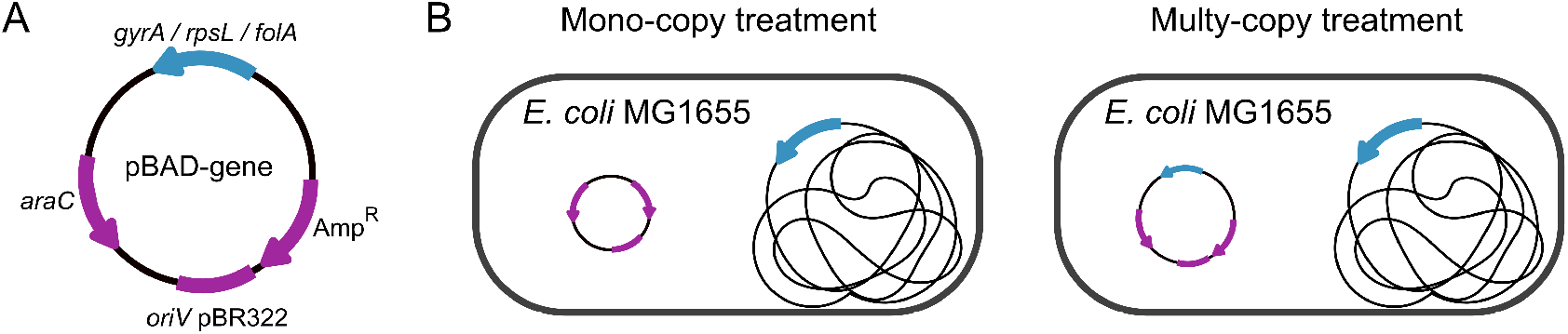
Construction of mono-copy and multi-copy experimental systems of housekeeping genes. (A) Schematic representation of plasmid pBAD carrying a housekeeping gene. (B) Construction of experimental system using *E. coli* MG1655 as recipient strain. To obtain completely isogenic backgrounds we transformed the empty pBAD vector in the mono-copy treatment strain. pBAD presents approximately 20 copies in the host bacterial cell.

**Figure S6.**
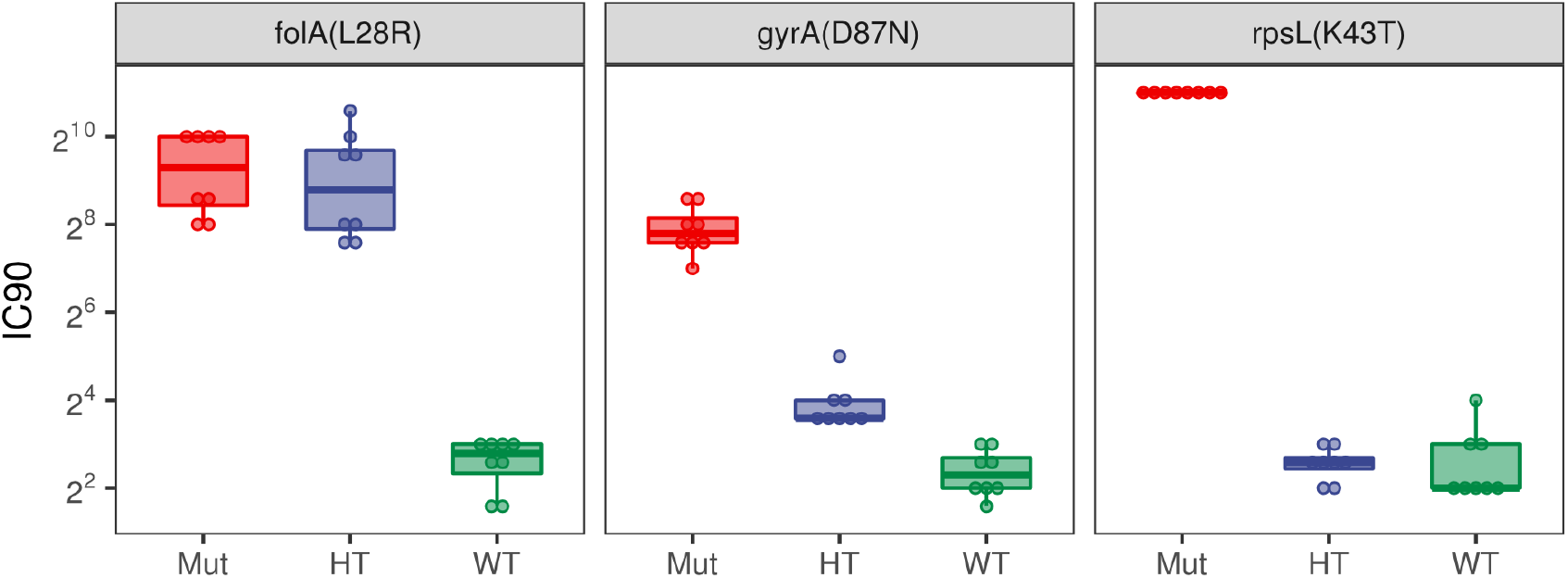
Antibiotic resistance levels of homozygous and heterozygous clones constructed to measure the coefficient of dominance of mutations. Antibiotic resistance phenotypes of the clones constructed to measure the coefficient of dominance of *folA*^L28R^ (trimethoprim), *gyrA*^D87N^ (nalidixic acid) and *rpsL*^K431^ (streptomycin) mutations. The inhibitory concentration 90 (IC_90_, in mg/L) of the homozygous mutant clones (Mut), heterozygous mutant clones (HT) and homozygous wild type clones (WT) are represented by boxes. The line inside the box marks the median. The upper and lower hinges correspond to the 25th and 75th percentiles. Note that for *folA*^L28R^ mutation the resistance level of HT clones is similar to that of the Mut clones (dominant mutations), while for *gyrA*^D87N^ and rpsL^K431^ mutations the resistance level of HT clones is similar to that of the WT clones (recessive mutations).

**Figure S7.**
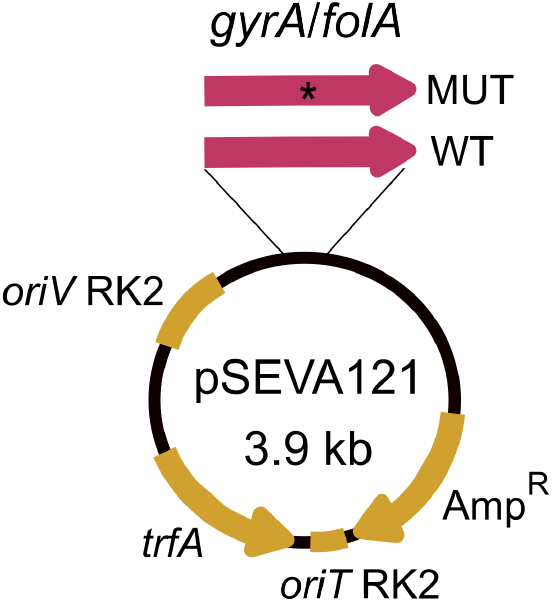
Construction of a mobilizable plasmid carrying *gyrA* or *folA* genes. Schematic representation of mobilizable plasmid pSEVA121. We cloned two different alleles of *gyrA* (WT and D87N) and *folA* (WT and L28R) into pSEVA121. pSEVA121 presents approximately 4 copies in the host bacterial cell.

**Figure S8.**
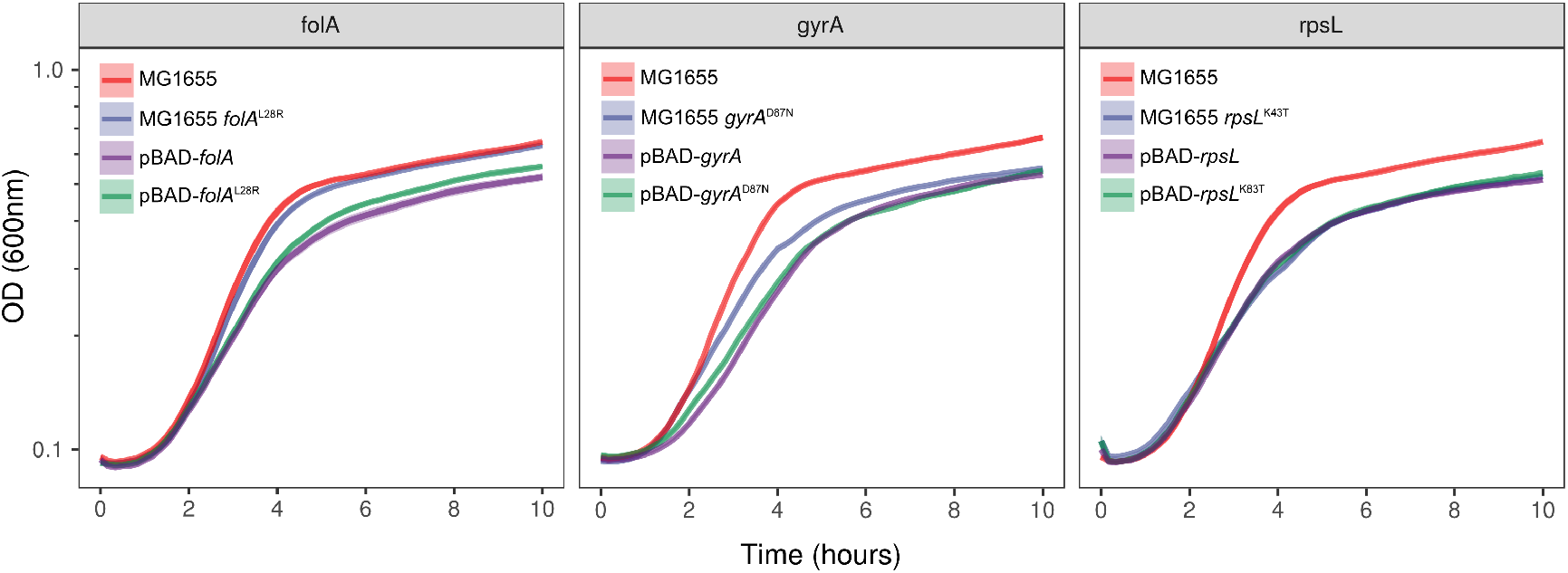
Growth curves of MG1655 with plasmid-and chromosomal-encoded wild type and mutated alleles of the genes under study. Growth curves measuring change in bacterial optical density at 600 nm (OD_600_, log scale) over time. Solid lines present the average of 6 biological replicates and the shaded region surrounding the curve represents standard deviation. Wild type *E. coli* MG1655 strain carrying an empty pBAD plasmid (red), MG1655 carrying the chromosomally encoded resistance-conferring mutant allele and an empty pBAD plasmid (blue), MG1655 carrying a pBAD plasmid encoding the wild type allele of the gene of interest (purple) and MG1655 carrying pBAD plasmid encoding a resistance-conferring mutant allele of the gene of interest (green). Note that none of the recessive resistance-conferring alleles (*gyrA*^D87N^ and *rpsL*^K43T^) present severe growth defects compared to the chromosomal mutants. Given that the frequency of wild-type strains carrying chromosomal mutant alleles is high in nature (Figure 4), our results suggest that the absence of mobile versions of these alleles is not due to the fitness costs imposed by the MGE-encoded alleles.

**Figure S9.**
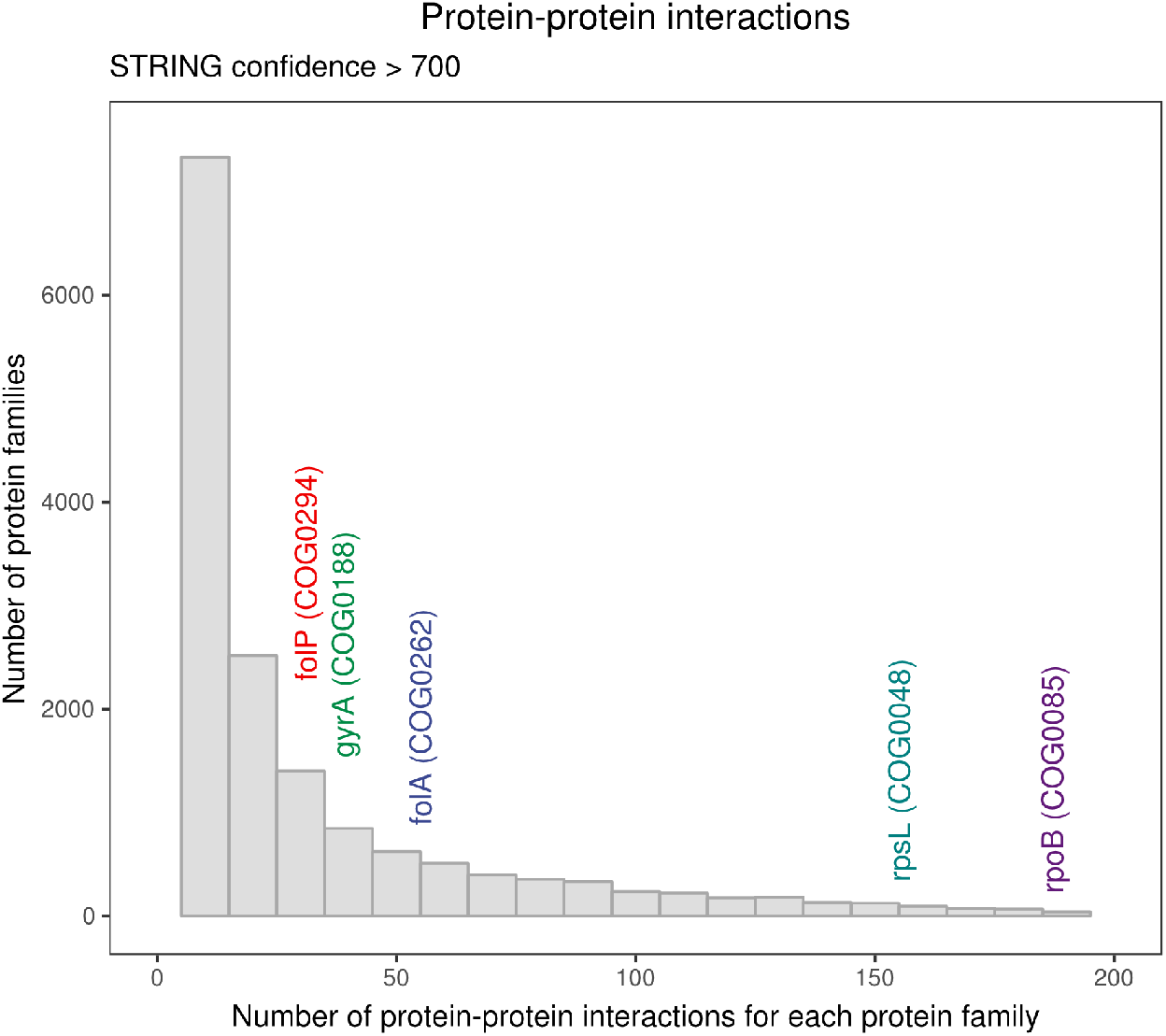
Connectivity of the proteins under study. Number of protein-protein interactions of the proteins under study according to STRING database (*41*). RpsL and RpoB are involved a high number of protein-protein interactions, which could constitute a barrier to their horizontal gene transfer (*4*). However, *gyrA*, which also confers antibiotic resistance through recessive mutations, encodes a protein with similar number of interactions as those conferring resistance through dominant mutations (e.g. *folA*), which are commonly present in MGEs (Figure 4).

**Figure S10.**
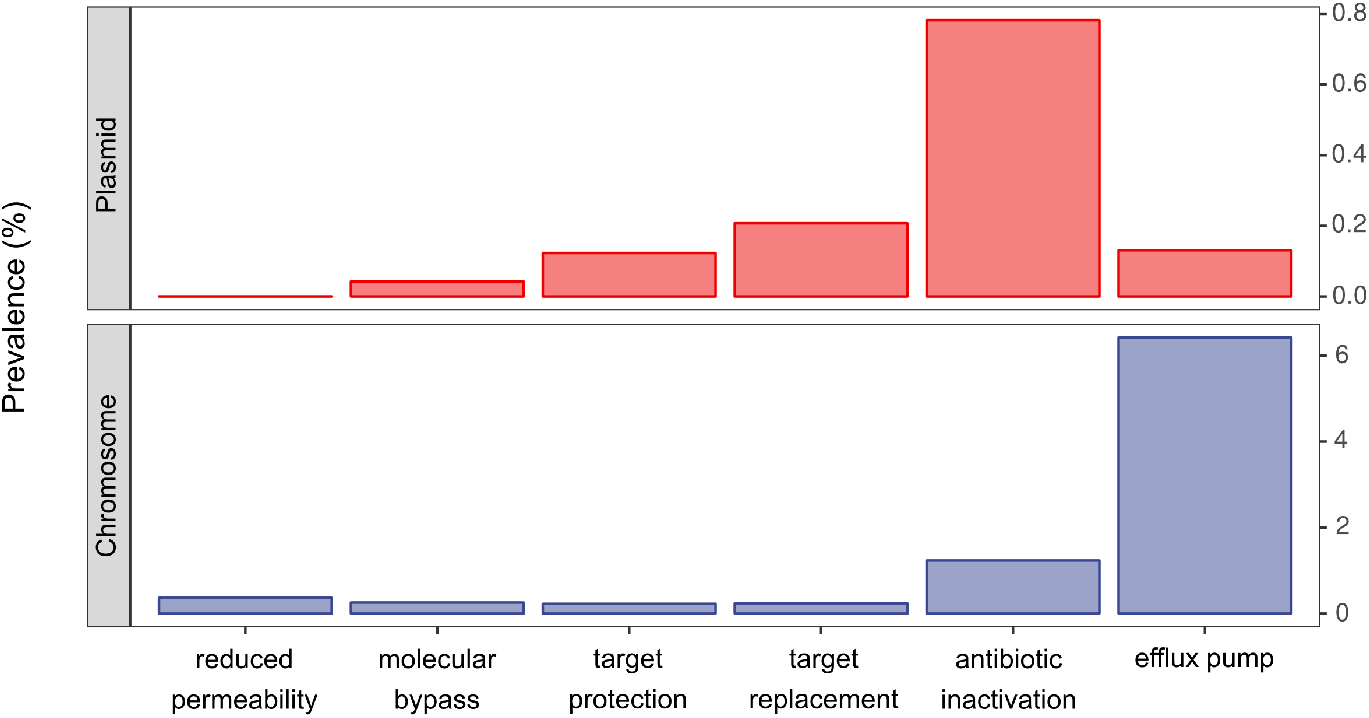
Distribution of antibiotic resistance genes in plasmids and chromosomes Frequency of genes belonging to the different categories of the Antibiotic Resistance Ontology group “mechanism of antibiotic resistance” in plasmids and chromosomes. We tried to explore the potential effect of genetic dominance on the general distribution of all resistance genes on plasmid/chromosomes according to their Antibiotic Resistance Ontology (ARO, grouped by “mechanism of antibiotic resistance”, data obtained from CARD). These groups include genes that are not necessary present on the chromosomes of recipient bacteria; therefore genetic dominance relationships between alleles may not impact their transferability. Actually, we hypothesized that those horizontally transferred genes with no pre-existing alleles on the recipient cells would be frequent in MGEs, since their phenotypic effect would not be masked regardless of the dominance of the allele. In line with this idea, we observed that the “antibiotic inactivation” category, which is mainly comprised of dedicated antibiotic degrading enzymes of xenogeneic origin, is extremely prevalent on plasmids. Conversely, categories including housekeeping genes vary in frequency on plasmids. Interestingly, a category enriched in housekeeping genes carrying loss of function (i.e. recessive) mutations is absent on plasmids. This is the category “reduced permeability to antibiotic”, which includes mostly inactivated outer membrane porins.

**Figure S11.**
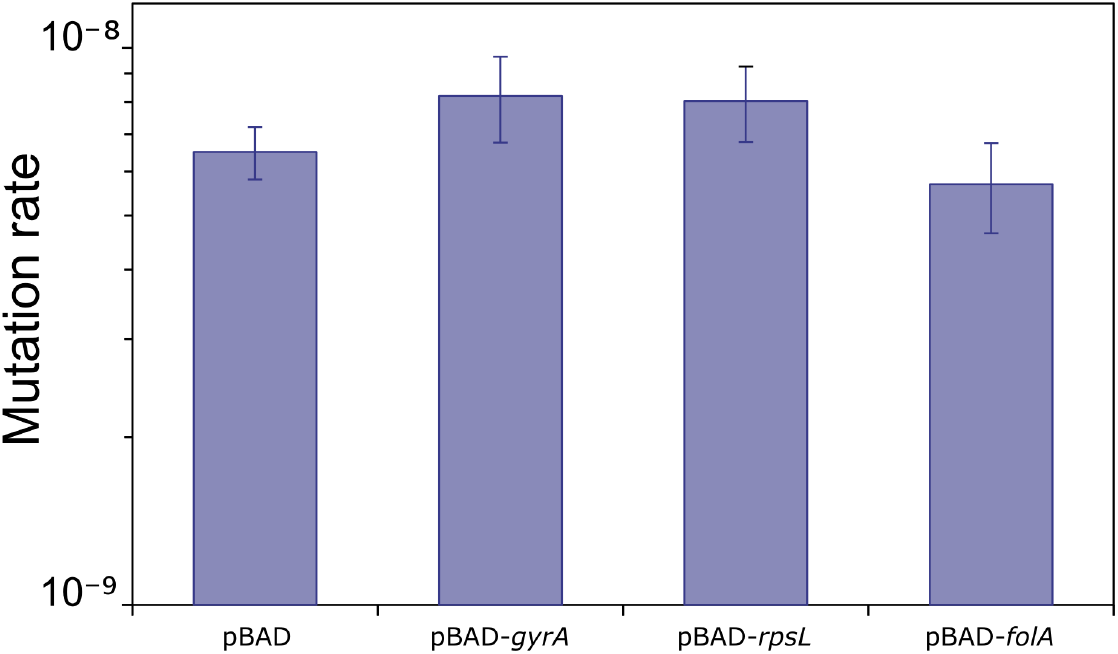
Rifampicin resistance mutation rates of the clones under study. Rifampicin resistance phenotypic mutation rates were performed to confirm the normo-mutator status of the clones used in this study. There were no significant differences in rifampicin mutation rates among strains (Likelihood Ratio Test P>0.13 in all cases).

## Supplementary Tables

Table S1: mutations described in the fluctuation assays.

Spreadsheet containing the mutations found after sequencing one clone of each biological replicate of the fluctuation assays, either from the mono-copy or the multicopy treatments (Figures 1C, 1D and 2C).

Table S2: Antibiotic susceptibility tests (IC_90_ results).

Spreadsheet containing the results of the IC_90_ assays used to calculate the coefficient of dominance in Figures 1D and 2C.

Table S3: Strains and plasmids used in this study.

Spreadsheet containing a short description of the strains and plasmids used in this study.

Table S4: Primers used in this study.

Spreadsheet containing the information about the oligonucleotides used in this study including their sequence, and a short description of their objective.

